# Characterization of the intrahippocampal kainic acid model in female mice with a special focus on seizure suppression by antiseizure drugs and DMSO

**DOI:** 10.1101/2022.07.05.498820

**Authors:** Melanie Widmann, Andreas Lieb, Angela Steck, Barbara Fogli, Anna Mutti, Christoph Schwarzer

**Author notes:** Corresponding author: Christoph Schwarzer, Institute of Pharmacology, Peter-Mayr-Straße 1a, 6020 Innsbruck, Austria.

## Abstract

**Objectives:** Affecting around 50 million people, men and women likewise, epilepsies are among the most common neurological diseases worldwide. Despite special challenges in the medical treatment of women with epilepsy, previous research has mainly focused on males, in particular preclinical animal studies, leaving a gap that needs to be urgently addressed. The intrahippocampal kainic acid (IHKA) mouse model of temporal lobe epilepsy (TLE) as one of the most frequently studied models in males is used for screening of novel antiepileptic therapies. In this study we investigate the IHKA model of TLE in female mice, in particular drug-resistance of hippocampal paroxysmal discharges. Furthermore, we provide evidence for anti-seizure effects of dimethyl sulfoxide (DMSO) in epileptic, but not naÏve mice.

**Methods:** After injecting KA unilaterally into the hippocampus of female mice, we monitored the development of epileptiform activity in *in-vivo* EEG recordings, evaluated responsiveness to the commonly prescribed antiseizure drugs (ASDs) lamotrigine (LTG), oxcarbazepine (OXC) and levetiracetam (LEV) and assessed typical neuropathological alterations of the hippocampus. Moreover, the effect of different doses of DMSO was tested in the IHKA chronic epilepsy model as well as on the PTZ-induced acute seizure threshold in both female and male mice.

**Results:** In the IHKA model, female mice replicated the key features of human TLE (EEG and neuropathological changes). Importantly, hippocampal paroxysmal discharges (HPDs) in female mice did not respond to commonly prescribed ASDs, thus representing a suitable model of drug-resistant seizures. The solvent DMSO caused a significant short-term reduction of HPDs, but did not affect the threshold of acute seizures.

**Significance:** By characterizing the drug-resistance of HPDs in the IHKA model of TLE in female mice we have laid a foundation for future research addressing sex-specific aspects. Considering the special issues complicating the therapeutic management of women, inclusion of females in the quest for novel treatment strategies is imperative. The observed effect of DMSO on epileptiform activity underlines that its application in epilepsy research is problematic and that the choice of solvent and appropriate vehicle control is crucial.

## INTRODUCTION

Affecting around 50 million people of both sexes and all ages, epilepsies are among the most common neurological diseases worldwide (Fiest et al., 2017) and contribute significantly to the global burden of diseases with an estimated proportion of more than 0.5% (Beghi et al., 2019). Although males and females display a similar prevalence, the medical management of women with epilepsy is particularly challenging due to complex interactions between sex hormones, seizure control and antiseizure drugs (ASDs) as well as special needs during pregnancy (for review see (Stephen et al., 2019; Taubøll et al., 2015)).

Accumulating evidence from both, preclinical research and clinical studies, indicates that female sex steroid hormones influence neuronal excitability: estrogens are generally regarded as eliciting excitatory effects primarily by potentiating glutamate responses, while progesterone and mainly its metabolite allopregnanolone (=3α-5α-tetrahydroprogesterone), exert an inhibitory effect via postsynaptic GABA_A_ receptors (Christian et al., 2020; Harden & Pennell, 2013; Herzog, 2008; Taubøll et al., 2015). Fluctuating hormone levels throughout the menstrual cycle can exacerbate seizure frequency during certain phases, which affects approximately one to two thirds of women with epilepsy (Bäckström, 1976; Herzog et al., 1997,2015; Laidlaw, 1956; Quigg et al., 2008). While women with epilepsy face special challenges, previous research, in particular preclinical animal studies mainly focused on males, leaving a gap that needs to be urgently addressed.

One of the most frequently studied models is the intrahippocampal kainic acid (IHKA) mouse model, which reproduces several EEG and histopathological features of human temporal lobe epilepsy (TLE) (Bouilleret et al., 2000; Riban et al., 2002; Suzuki et al., 1995). Hippocampal paroxysmal discharges (HPDs) were described as drug resistant seizures in males (Duveau et al., 2016; Riban et al., 2002; Zangrandi et al., 2016), however, there is still a remarkable lack of data on females. Recent studies suggest sex-specific differences regarding development and manifestation of seizures. IHKA in female mice resulted in faster onset of spontaneous seizures and no clear seizure-free latent period in contrast to males (Twele et al., 2016). In addition, seizure burden in females 2 months after IHKA was higher on pro/estrus compared with diestrus, characterized by increased time in seizures and longer seizure duration (Li et al., 2020). Importantly, drug-responsiveness of HPDs has not been studied in females so far.

Therefore, the first objective of this study was to further characterize the IHKA mouse model of TLE in female mice, in particular the drug-resistance of HPDs.

In the process of developing treatments for patients with drug-resistant seizures, poor water solubility is a frequent problem. Dimethyl sulfoxide (DMSO) is one of the most efficient solvents for water-insoluble drugs, however, its application in epilepsy research may be problematic due to a potential effect on neuronal excitability. First indications came from case reports about seizures in patients after stem cell transfusions containing DMSO as a cryoprotectant (Ataseven et al., 2017; Bauwens et al., 2005; Hequet et al., 2002; Maral et al., 2018; Martín-Henao et al., 2010; Mueller et al., 2007). Since data from *in vitro* and *in vivo* studies are inconsistent, reporting both pro-(Kovács et al., 2011; Kumari et al., 2018; Neuwelt et al., 1983; Tamagnini et al., 2014) and antiepileptic effects (Carletti et al., 2013; Kovács et al., 2011; Larsen et al., 1996; Maclennan et al., 1996; Tamagnini et al., 2014), we investigated the effects of DMSO on epileptiform activity in female and male IHKA mice and on the threshold of acute pentylenetetrazol (PTZ)-induced seizures.

## METHODS

### Animals

All experiments involving animals were approved by the Austrian Animal Testing Commission (Austrian Federal Ministry of Education, Science and Research) in accordance with Directive 2010/63/EU of the European Parliament and of the Council on the protection of animals used for scientific purposes. All experimental procedures were designed in compliance with the 3 Rs (replacement, reduction, refinement) principles.

Female and male adult C57BL/6N mice (2 - 12 months) were used, either bred in-house or purchased from Charles River (Sulzfeld, Germany). For breeding and maintenance, mice were group-housed (max. 5 mice in type II-L individually ventilated cages) under standard laboratory conditions (12 h light-dark cycle with lights on 7 am -7 pm, 22 ± 2 °C room temperature, 45 - 65 % relative humidity) with freely accessible food and water. Post-surgery mice were single-caged.

### Intrahippocampal kainic acid injection and electrode implantation

Stereotaxic surgery with IHKA injection and implantation of EEG electrodes was performed as previously described (Zangrandi et al., 2016). In short: Targeting the CA1 area of the left hippocampus (RC -1.8 mm, ML + 1.2 mm, DV -1.8 mm in reference to bregma), a bolus of 50 nl of 20 mM kainic acid (Ocean Produce International, Canada) solution was injected. After a waiting time of 10 min, the cannula was gradually retracted to minimize backflow along the injection track.

Following KA injection, 2 depth electrodes were implanted bilaterally into the CA1 area of the hippocampus (RC -1.8, ML ± 1.3, DV -1.8). Screw electrodes were secured above the motor cortex to record surface EEG and over the cerebellum as grounding and reference. An implant was built by fixing all electrodes and wires in place with Paladur® dental cement (Kulzer GmbH, Germany).

### EEG recordings and analysis

Wireless devices (Neurologger from either TSE systems, Germany or Evolocus, USA) simultaneously recording 4 channels (sampling rate 512 Hz or 500 Hz depending on manufacturer) were used for EEG recordings in freely moving animals. All recordings were performed in the home cages placed within a recording chamber covered by a mesh of conductive wire to block electromagnetic fields.

A custom-made python-based program was used for automated analysis. EEG traces were filtered between 0.5 and 70 Hz using a second-order Butterworth bandpass filter. Subsequently, the data were divided into 1 sec long intervals of which the minimum or maximum (depending on epileptic spike polarity) was determined and the mode calculated. The baseline for spike threshold detection was set to an amplitude minimum or maximum of twice the mode calculated above. Spike trains were defined as series of at least 3 epileptiform spikes (amplitude > 2x mode, width < 70 ms) lasting 1 – 10 sec with a minimum of 2 Hz. Trains of epileptiform spikes with a duration ≥ 10 sec were counted as HPDs. In addition, the EEG signals were visually scanned for generalized seizures using LabChart Reader version 8.1.14 (ADInstruments, New Zealand). Generalized seizures were defined as high-amplitude, high-frequency discharges simultaneously present in all channels followed by a clear post-ictal EEG depression. Using the fast Fourier transform (FFT) algorithm with a 10 sec sliding Hanning window the ipsilateral hippocampal EEG signal power was calculated, and divided into the following frequency bands which are typical for preclinical rodent EEG studies (Kadam et al., 2017): 1-4 Hz (delta), 4-8 Hz (theta), 8-13 Hz (alpha), 13-30 Hz (beta) and 30-80 Hz (gamma). In addition, the coastline, an aggregate of frequency and amplitude was computed (Lieb et al., 2018).

To evaluate the effect of treatments on epileptiform activity, the number and duration of spike trains and HPDs after the treatment were normalized to a baseline calculated from the 1h before the treatment. 5 min immediately before and after the injection were excluded from the analysis to minimize the confounding influence of handling stress. EEGs with < 100 sec spike trains and < 50 sec of HPDs, respectively, during the 1h pre-treatment baseline or a generalized seizure during the post-treatment period were excluded from the analysis.

### Nissl staining of frozen brain sections

After sacrificing mice by cervical dislocation, the brains were extracted and immediately snap-frozen in - 70°C 2-methylbutane. 20 µm coronal sections were cut on a cryotome (Microm Cryo-Star HM 560, Thermo Fisher Scientific, Germany) and mounted onto poly-L-lysine coated glass slides (Menzel GmbH & Co KG, Germany). Nissl staining was performed based on the protocol of (Paxinos & Watson, 1986). Briefly, after 10 min fixation in 4% PFA in 1x PBS the sections were dehydrated in a graded ethanol series (70%, 95%, 100%, 100%) and transferred to butyl acetate. Rehydration in a reverse ethanol series was followed by incubation in 0.5% cresyl violet solution for 20 min. Subsequently, the sections were again dehydrated in an increasing ethanol series and cleared in butyl acetate. Finally, the sections were embedded with Eukitt® mounting medium (ORSAtec GmbH, Germany) and covered with glass coverslips. Images were taken with a Zeiss Axiophot microscope with a 2.5x/0.075 Plan-Neofluar objective connected to a AxioCam MRc5 camera (Carl Zeiss AG, Germany).

### Pentylenetetrazol seizure threshold

To determine the threshold for PTZ-induced acute seizures, PTZ (10 mg/ml) was infused at a rate of 100 µl/min into the tail vein of freely moving animals. Upon onset of generalized seizures, the infusion was stopped and the animal sacrificed by cervical dislocation. The seizure threshold dose (in mg/kg PTZ) was calculated from the infused volume of PTZ solution in relation to the body weight as described in (Loacker et al., 2007).

### Drugs and solutions

PTZ (Merck KGaA, Germany) was dissolved in 0.9 % saline. 20 mM kainic acid (Ocean Produce International, Canada) solution was prepared in purified water. LTG, OXC, LEV (all from BioTechne, USA) and diazepam (Takeda Austria GmbH, Austria) were dissolved in vehicle containing 90 % saline and 10 % DMSO with 3 % Tween 80. Fresh DMSO was used either pure 100 % or diluted in 0.9 % saline with concentrations ranging from 1 % to 30 %. Lidocaine (BioTechne, USA) was solubilized in either 100 % fresh DMSO or 0.9 % saline.

### Data and statistical analysis

For statistical analysis GraphPad Prism version 9.3.1 for macOS (GraphPad Software, USA) was used. The development of epileptiform activity in female mice was analyzed by fitting a 1-way mixed-effects model for repeated measures and a post test for linear trend. In the ASDs experiment, the pre-/post-treatment ratio of different parameters was analyzed with a 1-way mixed-effects model for repeated measures followed by Dunnett’s multiple comparisons test. The occurrence of generalized seizures after ASDs treatments was analyzed with Chi-square test. Repeated measures 1-way ANOVA followed by Dunnett’s multiple comparisons test was used for analyzing power bands in the ASDs experiment. In the DMSO experiment, 2-way mixed-effects model for repeated measures followed by Dunnett’s multiple comparisons test was applied. The experiment with DMSO and lidocaine was analyzed with 1-way mixed-effects model for repeated measures followed by Šídák’s multiple comparisons test to compare lidocaine-saline with the saline control group and lidocaine-DMSO with the DMSO group. Data from the PTZ seizure threshold determination were analyzed using ordinary 2-way ANOVA. P-values smaller than 0.05 were considered statistically significant. Data are presented as mean ± standard deviation (SD), with data for individual animals also shown.

## RESULTS

### Post-KA Development of Epileptiform Activity in Female Mice

To evaluate the development of epileptiform activity, 18 female mice after IHKA injection were monitored with weekly recordings during the first 5 weeks and at 2 months. In general, female mice presented the same type of epileptiform activity, namely spike trains, HPDs and generalized seizures (Figure 1 A and B), as previously described in male IHKA mice. Based on the manifestation of HPDs at 2 months 2 subgroups (>50 sec/h HPDs termed high-HPDs group, <50 sec/h HPDs termed low-HPDs group) were analyzed in detail. This grouping of animals took into account that mice with >50 sec/h HPDs are suitable to test potential antiepileptic interventions, because they are considered as having stably established chronic focal epilepsy.

**Figure 1.**
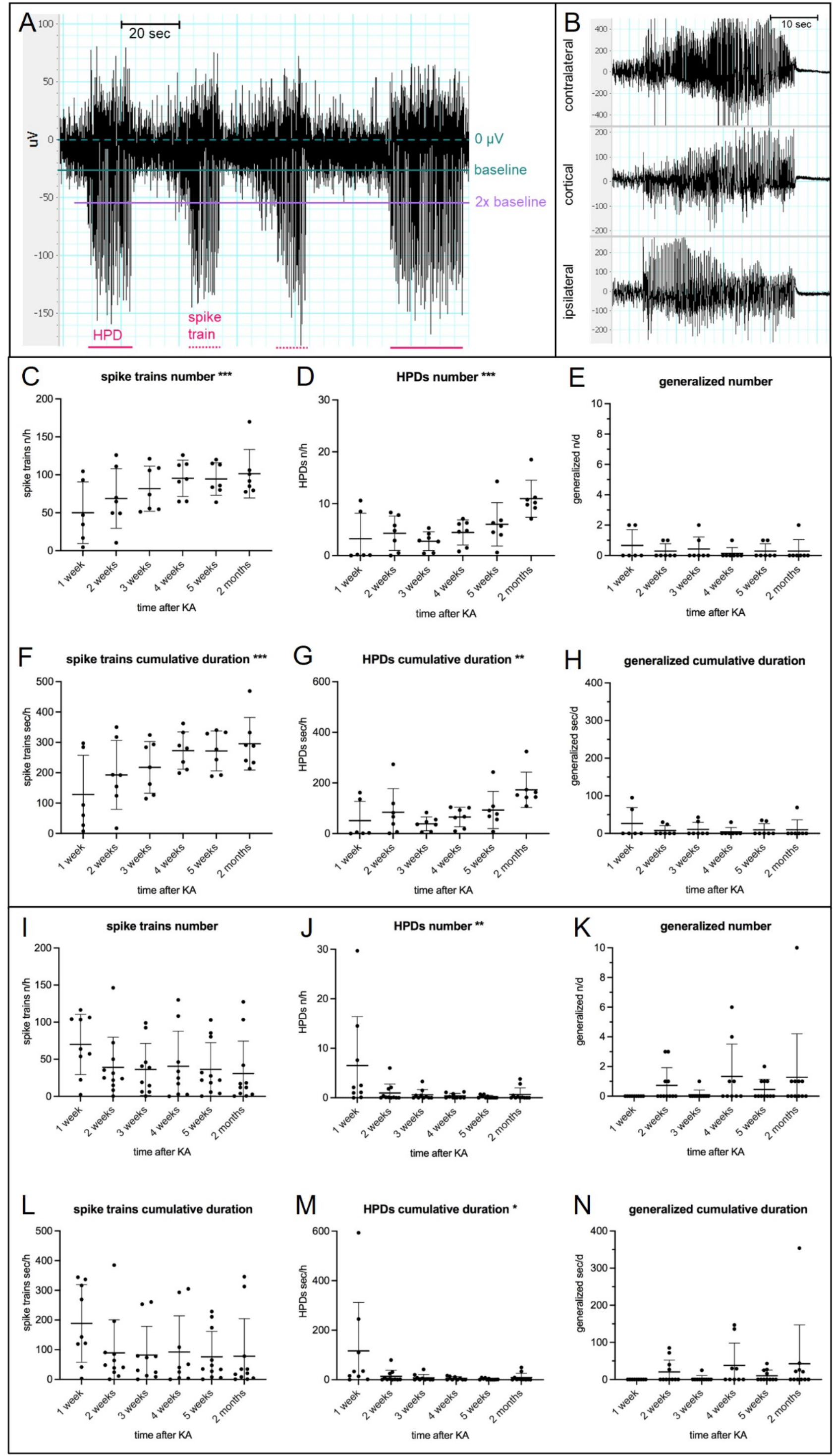
Development of epileptiform activity in female mice after IHKA. (A) Representative EEG trace of the ipsilateral hippocampus showing spike trains (dotted magenta lines) and HPDs (continuous magenta lines). Spike trains are series of at least 3 epileptiform spikes with a duration ≥1 and <10 sec and a maximum interspike interval of 1.5 sec. Trains of epileptiform spikes lasting ≥10 sec are counted as HPDs. In this exemplary trace the spike direction is negative, a minority of animals, however, present with positive spikes. Dashed petrol line = 0 µV; continuous petrol line = baseline calculated automatically based on the mode of the amplitude minima; lilac line = 2x baseline defining threshold for epileptiform spikes. (B) EEG traces of a typical generalized seizure with approximately 30 sec of high-amplitude, high-frequency discharges simultaneously both in the ipsi- and contralateral hippocampus as well as in the cortical electrode followed by a pronounced post-ictal depression of EEG activity. (C-H) Development of epileptiform activity in the female high-HPDs group (>50 sec/h HPDs at 2 months, n=7) showing increasing (C) number [n/h] and (F) cumulative duration [sec/h] of spike trains and increasing (D) number [n/h] and (G) cumulative duration [sec/h] of HPDs. (E) Number [n/d] and (H) cumulative duration [sec/d] of generalized seizures. (I-N) Females of the low-HPDs group (<50 sec/h HPDs at 2 months, n=11) showing a more heterogenous development of (I) number [n/h] and (L) cumulative duration [sec/h] of spike trains and decreasing (J) number [n/h] and (M) cumulative duration [sec/h] of HPDs. (K) Number [n/d] and (N) cumulative duration [sec/d] of generalized seizures. Note one animal with strikingly higher seizure activity increasing up to 10 generalized seizures at the 2 months recording time point. Data in the scatter plot are presented as mean ± SD. One-way mixed-effects model for repeated measures followed by a post test for linear trend (significance levels shown in graph). *\* p = 0*.*01-0*.*05;* ** p = 0.001-0.01; *** p = 0.0001-0.001

Females of the high-HPDs group (n=7) significantly increased spike trains (Figure 1C and F) and HPDs (Figure1D and G) during the observation period up to 2 months after IHKA [one-way mixed-effects model for repeated measures followed by post test for linear trend: spike trains n/h: F (1, 29) = 17.55, p=0.0002; spike trains sec/h: F (1, 29) = 18.45, p=0.0002; HPDs n/h: F (1, 35) = 16.85, p=0.0002; HPDs sec/h: F (1, 35) = 9.623, p=0.0038]. Regarding generalized seizures (Figure 1 E and H), there were no significant differences between the different time points. 3 female mice (43%) had exclusively focal epileptic activity without generalized seizures, while 4 (57%) developed at least 1 generalized seizure during the whole observation period. Mice of the high-HPDs group showed a maximum of 2 generalized seizures per day.

Female mice displaying <50 sec/h HPDs at 2 months (n=11), in contrast, showed a more heterogenous development of epileptiform activity (Figure 1I and L for spike trains, J and M for HPDs, K and N for generalized) with a significant decrease of HPDs over time [number of HPDs: F (1, 55) = 8.042, p=0.0064; cumulative duration of HPDs: F (1, 55) = 6.990, p=0.0107]. While 4 females (36%) did not show any generalized seizures and are considered as not having developed epilepsy, 7 (64%) had at least one generalized seizure throughout the recording period and are therefore classified as epileptic. One female of the low-HPDs group showed strikingly higher generalized seizure activity, increasing even up to 10 generalized seizures during 24 h at the 2 months recording time point.

### Effect of ASDs in Female IHKA Mice

To evaluate the effect of frequently prescribed new generation ASDs, 7 female IHKA mice were tested. Lamotrigine (LTG; 10 and 30 mg/kg), oxcarbazepine (OXC; 10 and 30 mg/kg), levetiracetam (LEV; 100 and 300 mg/kg), diazepam (DZP; 2.5 mg/kg) as positive control and vehicle (10% DMSO and 90% saline with 3% Tween 80) were administered i.p. in a crossover design (injection volume 10 µl/g). Number and cumulative duration of spike trains and HPDs were assessed in the 35-94 min after treatment to account for the delayed onset of the ASDs effect and normalized to a baseline period of the last hour before the treatment (Figure 2).

**Figure 2.**
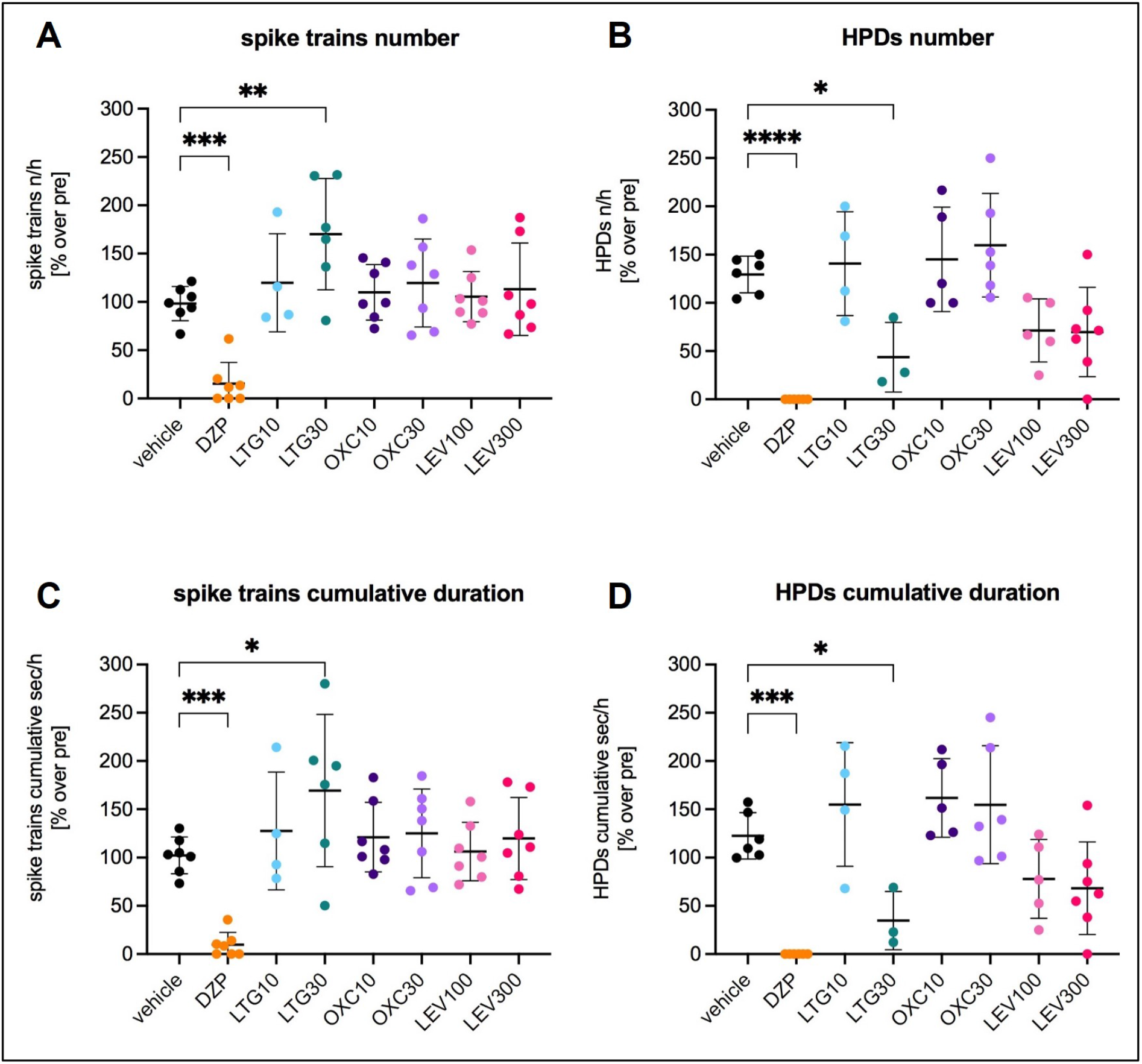
Effect of frequently prescribed new generation ASDs in female IHKA mice. (A) Number [% over pre] and (C) cumulative duration [% over pre] of spike trains and (B) number [% over pre] and (D) cumulative duration [% over pre] of HPDs 35 – 94 min after treatment. Among the tested ASDs only the higher dose of LTG caused a significant reduction of HPDs (number and cumulative duration), while simultaneously increasing spike trains (number and cumulative duration), corresponding to a reduced duration of epileptiform events. The benzodiazepine DZP highly significantly reduced both spike trains and HPDs. One-way linear mixed model for repeated measures followed by Dunnett’s multiple comparisons test. Data (n=7) are presented as % normalized to the pretreatment period (mean ± SD). DZP, diazepam; LTG, lamotrigine; OXC, oxcarbazepine; LEV, levetiracetam. * p = 0.01-0.05; ** p = 0.001-0.01; *** p = 0.0001-0.001; **** p = < 0.0001

Among the tested ASDs only the higher dose of LTG (30 mg/kg) caused a statistically significant reduction of number (Figure 2B; 43.7 ± 36.1, one-way linear mixed model for repeated measures followed by Dunnett’s multiple comparisons test 95% CI 6.025 to 165.5, p=0.0304) and cumulative duration of HPDs (Figure 2 D; 34.7 ± 30.2, 95% CI 2.963 to 172.8, p=0.0401) in comparison to vehicle (number of HPDs 129.4 ± 19.1, cumulative duration of HPDs 122.5 ± 24.1). The reduction of HPDs, however, was paralleled by a significant increase of number (Figure 2 A; 170.2 ± 57.7, 95% CI -124.8 to -21.48, p=0.0025) and cumulative duration of spike trains (Figure 2 C; 169.4 ± 78.8, 95% CI -126.1 to -10.70, p=0.0140) after LTG 30 mg/kg compared to vehicle (number of spike trains 98.3 ± 17.7; cumulative duration of spike trains 102.3 ± 18.9). This shift from HPDs towards spike trains corresponds to a decrease of the duration of epileptiform events after LTG 30 mg/kg. Intriguingly, we observed a significant clustering of generalized seizures in the 94 min after LTG (3 animals after 10 mg/kg and 1 after 30 mg/kg; Chi-square p=0.0160), whereas none of the animals developed generalized seizures after the other treatments. As expected, DZP resulted in a highly significant reduction of both number (15.3 ± 22.0, 95% CI 33.58 to 132.4, p=0.0003) and cumulative duration of spike trains (9.8 ± 12.8, 95% CI 37.29 to 147.7, p=0.0003) in comparison to vehicle (number of spike trains 98.3 ± 17.7; cumulative duration of spike trains 102.3 ± 18.9). HPDs (number 0.0 ± 0.0, 95% CI 64.32 to 194.5, p=<0.0001, cumulative duration 0.0 ± 0.0, 95% CI 53.23 to 191.7, p=0.0002) were completely eliminated after DZP compared to vehicle (number of HPDs 129.4 ± 19.1, cumulative duration of HPDs 122.5 ± 24.1).

In the power spectrum (Figure 3 D) LTG 30 mg/kg showed in the 35-94 min a pronounced increase in the delta frequency range compared to the pretreatment period. In accordance, power in the 1-4 Hz frequency band (35-94 min after treatment normalized to the pretreatment period; Figure 4 A) increased statistically significant after LTG 30 mg/kg (143.5 ± 44.5, repeated measures one-way ANOVA followed by Dunnett’s multiple comparisons test 95% CI -65.72 to -8.620, p=0.0059) in comparison to vehicle (106.3 ± 14.3). DZP (Figure 3 B) caused a marked reduction over the whole power spectrum. Accordingly, the 1-4 Hz (33.2 ± 8.8, 95% CI 44.56 to 101.7, p=<0.0001; Figure 4 A), 4-8 Hz (36.0 ± 9.3, 95 % CI 47.97 to 89.83, p=<0.0001; Figure 4 B), 8-13 Hz (50.4 ± 14.8, 95 % CI 30.32 to 76.03, p=<0.0001; Figure 4 C) and 13-30 Hz (72.9 ± 16.0, 95% CI 9.219 to 53.53, p=0.0024; Figure 4 D) frequency bands revealed a statistically significant reduction in comparison to vehicle (1-4 Hz: 106.3 ± 14.3; 4-8 Hz: 104.9 ± 6.7; 8-13 Hz: 103.5 ± 9.4; 13-30 Hz: 104.3 ± 10.6). Also the coastline (Figure 4 F), an aggregate of amplitude and frequency, was significantly reduced after DZP (85.0 ± 13.9, 95% CI 3.410 to 48.08, p=0.0177) compared to vehicle (110.7 ± 14.0).

**Figure 3.**
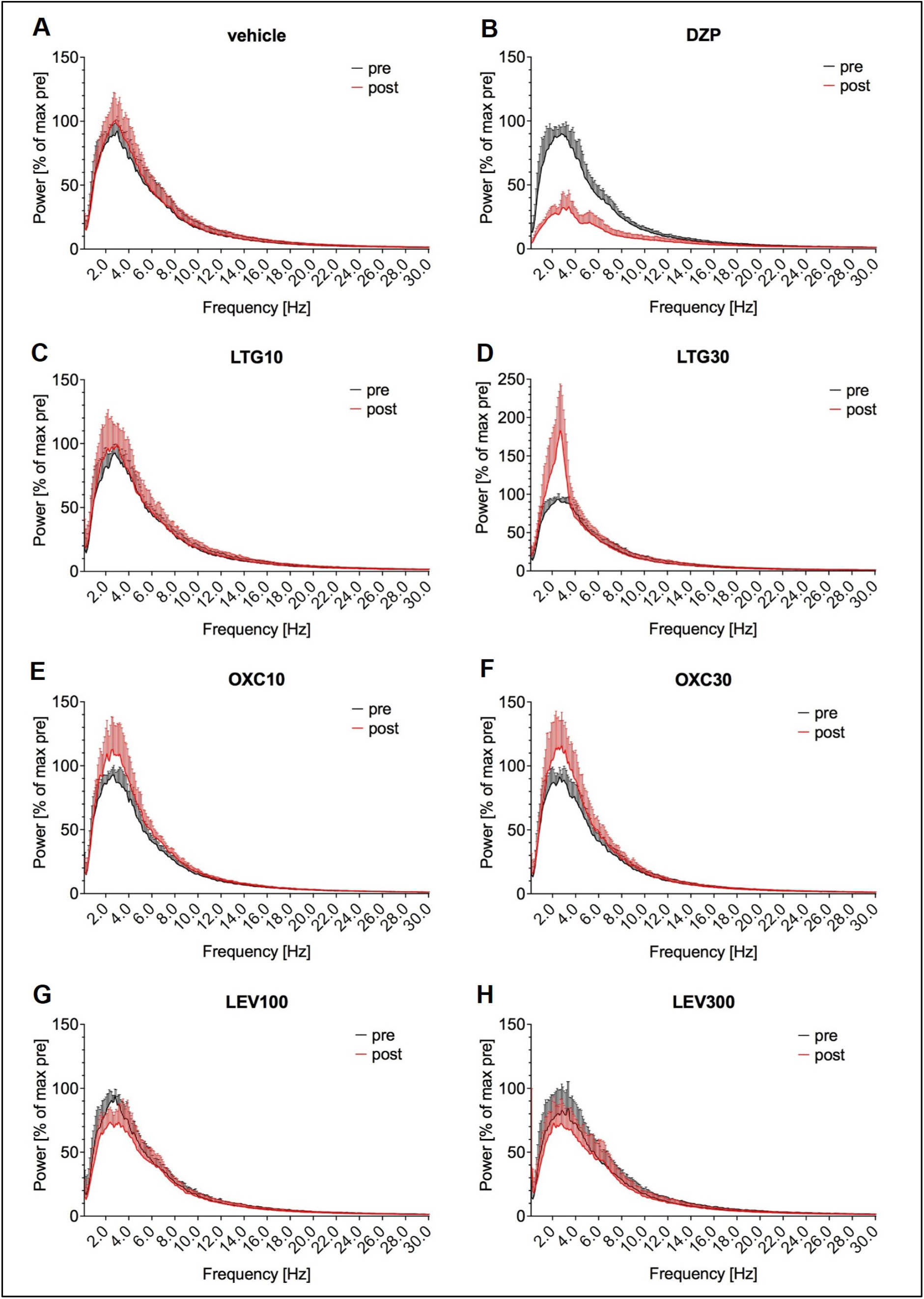
Power spectral analysis of the effect of frequently prescribed new generation ASDs in female IHKA mice. The periodogram shows after DZP (B) a marked decrease over the whole power spectrum and after LTG30 (D) a pronounced increase in the delta frequency range. Note the different y-axis range for LTG30. Power in µV^2^/Hz in 5 sec was calculated for the 64-5 min before (black) and 35-94 min after the treatment (red), normalized to the maximum of the pretreatment period and plotted as mean ± SD [% of pretreatment maximum].

**Figure 4.**
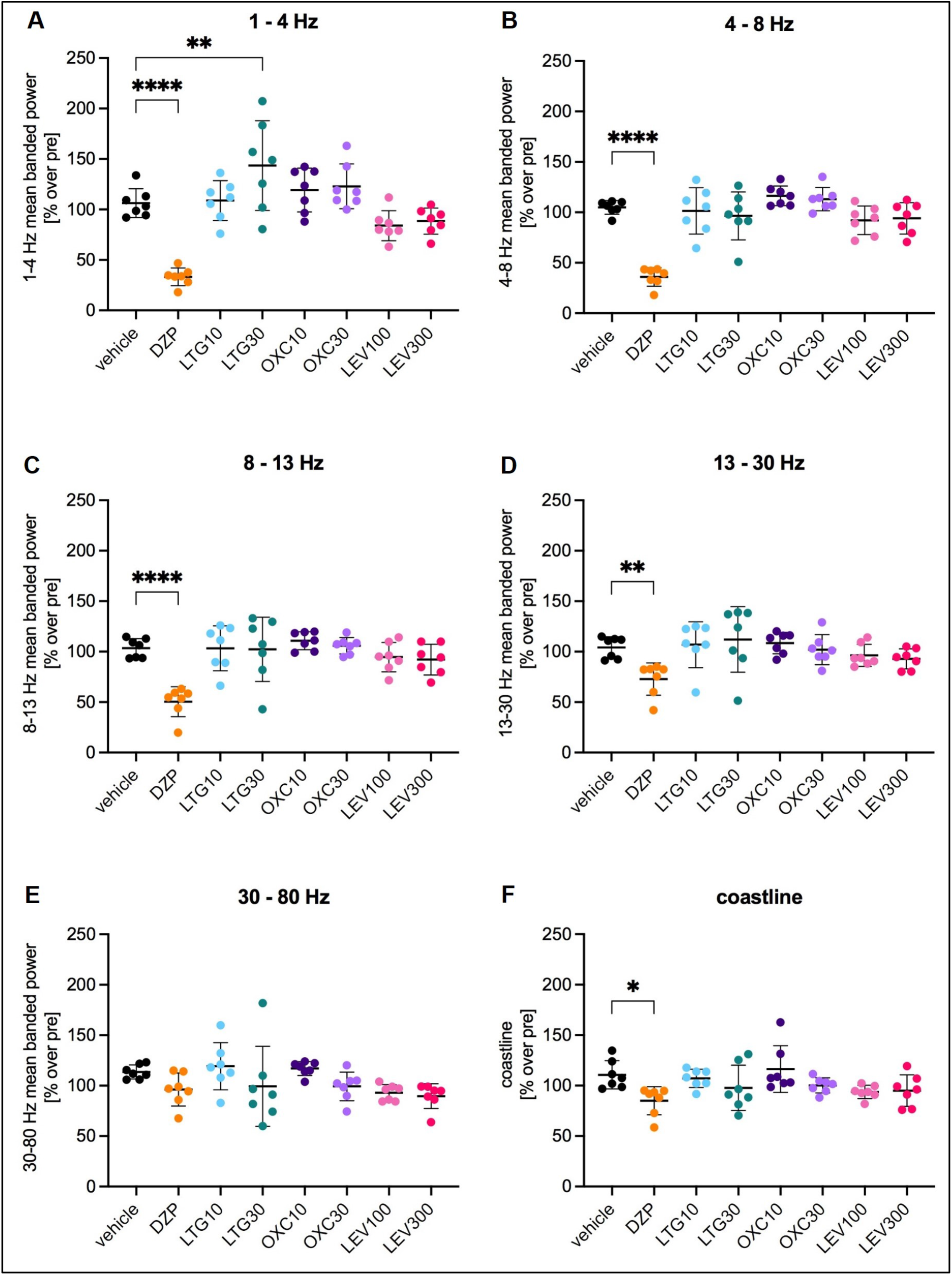
Banded power after widely prescribed new generation ASDs in the female IHKA mouse model of TLE. (A) Power in the 1-4 Hz, (B) 4-8 Hz, (C) 8-13 Hz, (D) 13-30 Hz and (E) 30-80 Hz frequency bands as well as (F) the coastline were calculated for the 35-94 min after treatment normalized to the pretreatment period. LTG30 caused a significant increase in the 1-4 Hz band, DZP significantly decreased the 1-4 Hz, 4-8 Hz, 8-13 Hz and 13- 30 Hz bands and the coastline. Data (n=7) are presented as % normalized to the pretreatment period (mean ± SD) and were analyzed with a repeated measures one-way ANOVA followed by Dunnett’s multiple comparisons test. DZP, diazepam; LTG, lamotrigine; OXC, oxcarbazepine; LEV, levetiracetam. * p = 0.01-0.05; ** p = 0.001- 0.01; **** p = < 0.0001

### Morphological Evaluation of Female IHKA Mice

Nissl-stained brain sections (n=18) were analyzed after the experimental series with ASDs to assess morphological alterations related to hippocampal sclerosis (Figure 5). For this morphological analysis, mice from the high-HPDs group and mice with generalized seizures from the low-HPDs group were summarized as epileptic group (n=14). The remaining animals from the low-HPDs group with neither sufficient HPDs nor generalized seizures were categorized as non-epileptic (n=4). All epileptic mice showed typical neuropathological changes of the KA-injected hippocampus. 57% (n=8) displayed extensive loss of Nissl-stained cells in CA1 and CA3 accompanied by massive dispersion of the granule cells. 43% (n=6) showed CA1 cell loss combined with granule cell dispersion, while the CA3 subregion appeared mainly unaffected. 2 of the 4 non-epileptic mice were excluded from the analysis because they had been reinjected with KA. Of the remaining 2 non-epileptic animals, one did not develop any of the typical morphological alterations, whereas the other one showed CA1 and CA3 cell loss as well as granule cell dispersion.

**Figure 5.**
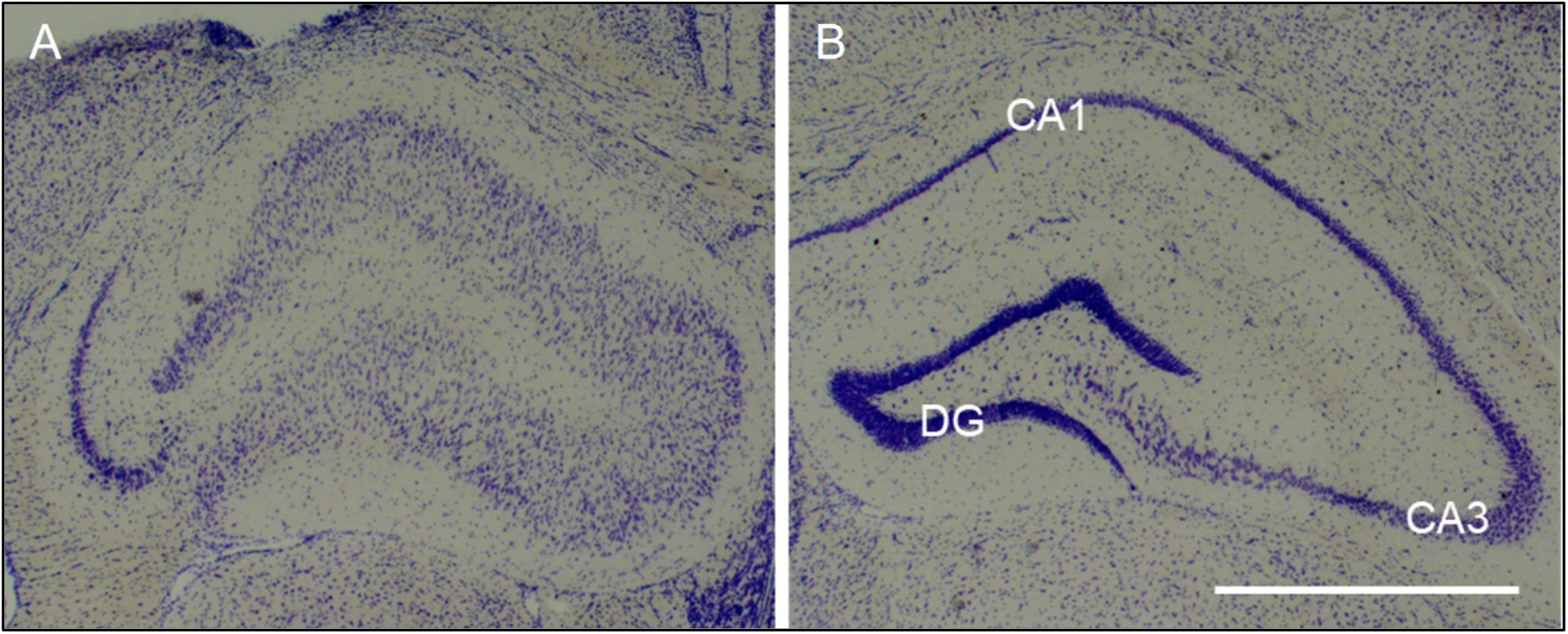
Ipsilateral neuropathological changes after IHKA in female mice. (A) Representative microscope image showing the characteristic pattern of ipsilateral morphological alterations with disintegration of the normal hippocampal layer structure, characterized by extensive neuronal loss in CA1 and less pronounced in CA3 as well as dispersion of granule cells, in a 20 µm Nissl-stained section of the dorsal hippocampus near the injection site (RC -1.8 mm). The tissue damage above the hippocampus is caused by removal of the depth electrode. (B) In the contralateral non-injected hippocampus layer integrity is mostly preserved. CA, cornu ammonis; DG, dentate gyrus; Scale bar = 1 mm

### Effect of DMSO in the IHKA Mouse Model of TLE

Using female (n=9) and male (n=8) IHKA mice, we evaluated the effect of different concentrations of DMSO (1%, 10%, 30% and 100% DMSO corresponding to 16.5 mg/kg, 165.1 mg/kg, 495.2 mg/kg and 1651 mg/kg vs. saline s.c. 1.5 µl/g) on epileptiform EEG activity (Figure 6 A-D). Number and cumulative duration of spike trains and HPDs were analyzed in the 5-34 min after treatment normalized to a baseline of the last hour before the treatment. While DMSO administration had no statistically significant effects on spike trains in female mice, in male mice the cumulative duration of spike trains (Figure 6 C) showed a significant reduction after 100% DMSO (75.6 ± 45.3, two-way linear mixed model for repeated measures followed by Dunnett’s multiple comparisons test 95% CI 7.221 to 109.4, p=0.0204) compared to saline (133.9 ± 20.7). Regarding HPDs (Figure 6 B and D), both females and males displayed a significantly reduced number (females 40.8 ± 47.1; 95% CI 12.31 to 147.9, p=0.0159; males 19.2 ± 23.1; 95% CI 15.79 to 142.9, p=0.0100) and cumulative duration of HPDs (females 37.3 ± 42.5; 95% CI 21.46 to 155.9, p=0.0062; males 16.9 ± 19.0; 95% CI 13.63 to 139.6, p=0.0125) after 100% DMSO in comparison to saline (number of HPDs: females 120.9 ± 34.8, males 98.5 ± 64.9; cumulative duration of HPDs: females 126.0 ± 43; males 93.6 ± 58.0). Lower DMSO concentrations (1%, 10% and 30%) did not cause significant changes of spike trains or HPDs compared to saline.

**Figure 6.**
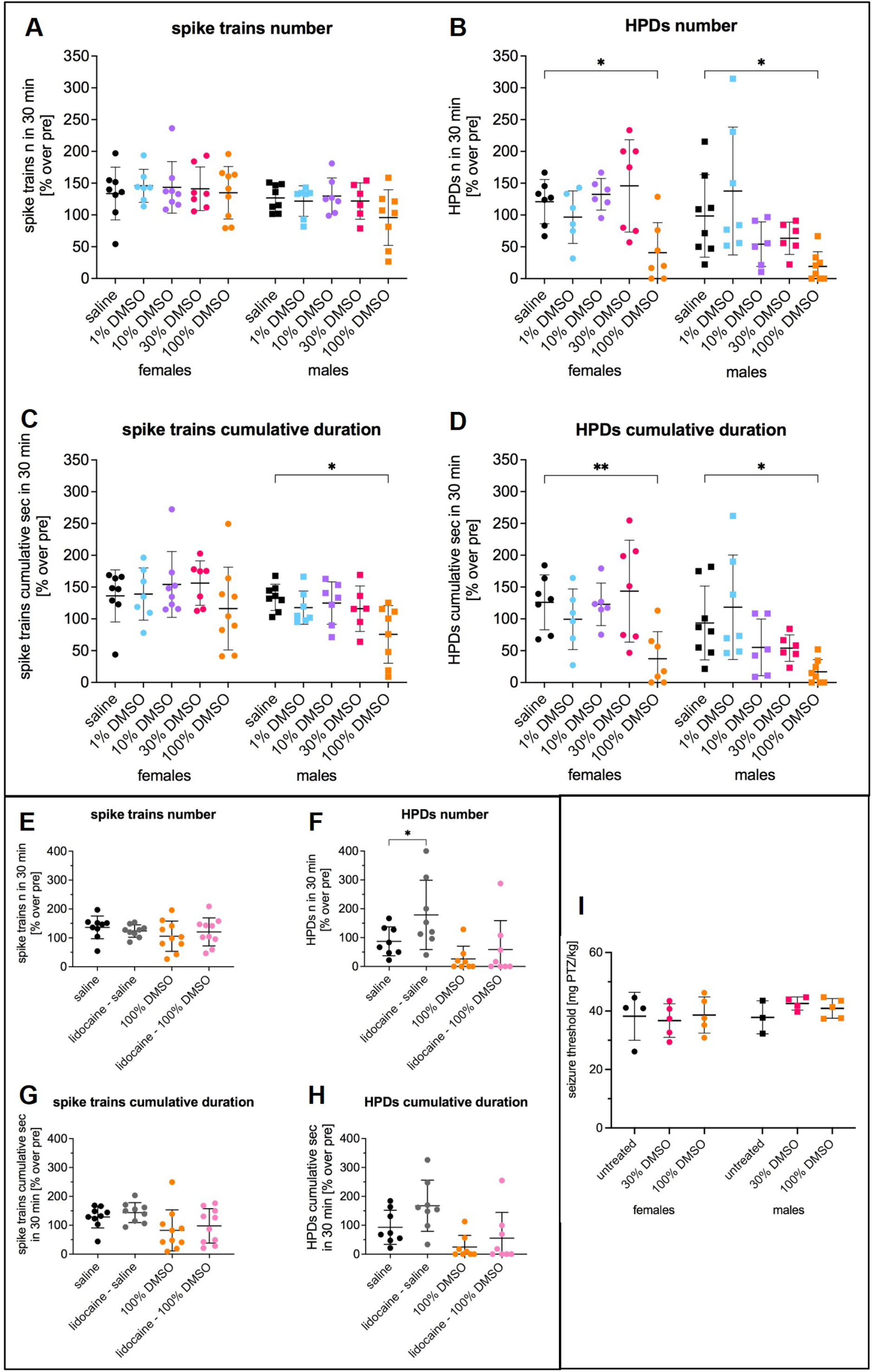
Effect of different concentrations of DMSO in IHKA mice and on PTZ-induced acute seizure threshold. (A-D) Effect of 1%, 10%, 30% and 100% DMSO compared to saline in female and male IHKA mice on (A) number [% over pre] and (C) cumulative duration [% over pre] of spike trains and (B) number [% over pre] and (D) cumulative duration [% over pre] of HPDs in 5-34 min after treatment. 100% DMSO in male mice resulted in a significant reduction of the cumulative duration of spike trains compared to saline. Regarding HPDs, number and cumulative duration were significantly reduced after 100% DMSO in both females and males in comparison to saline. Data (females n=9, males n=8) are presented as % normalized to the pretreatment period (mean ± SD) and were analyzed with a 2-way linear mixed model for repeated measures followed by Dunnett’s multiple comparisons test. * p = 0.01-0.05; ** p = 0.001-0.01 (E-H) Effect of lidocaine dissolved in saline and lidocaine dissolved in 100% DMSO compared to the respective control group in IHKA mice on (E) number [% over pre] and (C) cumulative duration [% over pre] of spike trains and (B) number and (D) cumulative duration of HPDs in 5-34 min after treatment. Addition of lidocaine to DMSO did not have a statistically significant effect compared to DMSO alone. Lidocaine dissolved in saline significantly increased the number of HPDs compared to the saline control group. Pooled data from females (n=7) and males (n=3) are presented as % normalized to the pretreatment period (mean ± SD) and were analyzed with a 1-way linear mixed model for repeated measures followed by Šídák’s multiple comparisons test. * p = 0.01-0.05 (I) Seizure threshold [mg PTZ/kg] determined by PTZ tail-vein infusion in untreated animals or 30 min after 30% or 100% DMSO showed no statistically significant differences. Data (n=3–5) are expressed as mean ± SD and were analyzed with ordinary 2-way ANOVA.

In addition, to exclude a potential confounding influence of local painful sensations after DMSO injection, we treated a subgroup (females n=7, males n=3) with DMSO combined with the local anesthetic lidocaine 4 mg/kg (Figure 6E-H). Lidocaine, if administered systemically, could influence epileptic activity via its effect on voltage-gated sodium channels. Although serious systemic effects are unlikely with s.c. lidocaine injections in the recommended concentration for local anesthesia, we additionally tested lidocaine dissolved in saline for potential effects on epileptiform activity. Addition of lidocaine to DMSO did not show a statistically significant effect in any of the analyzed parameters in comparison to DMSO alone suggesting that a potential painful sensation is not involved in the observed antiepileptic effect of DMSO. Intriguingly however, lidocaine in saline significantly increased the number of HPDs (Figure 6F; 178.9 ± 120.1; one-way linear mixed model for repeated measures followed by Šídák’s multiple comparisons test 95% CI -182.7 to -0.8838, p=0.0476) in comparison to the saline control group (87.1 ± 50.3).

### Lack of Effect of DMSO on Acute Seizures (PTZ seizure threshold)

To evaluate whether DMSO also influences acute seizures in non-epileptic mice (females n=14, males n=12), we determined the seizure threshold by infusing the GABA_A_ receptor antagonist PTZ via the tail vein in untreated mice or 28 min after 30% (495 mg/kg) or 100% DMSO (1651 mg/kg) 1.5 µl/g s.c. (Figure 6I). Statistical analysis using ordinary two-way ANOVA showed no significant differences, suggesting that DMSO does not affect the acute seizure threshold.

## DISCUSSION

In the first part of this study, we characterized the IHKA model in female mice regarding development of seizures, hippocampal neuropathological alterations and in particular responsiveness of HPDs to ASDs.

Within 2 months after IHKA, female mice developed epileptiform EEG activity (spike trains, HPDs and generalized seizures) as we had previously observed in male IHKA mice (Zangrandi et al., 2016). It is particularly noteworthy that female IHKA mice in our study established HPDs, which were validated as a model of drug resistant seizures in male mice (Duveau et al., 2016; Riban et al., 2002; Zangrandi et al., 2016). This contrasts with the findings of Twele *et al*. describing HPDs frequently in males, but almost never in females of various mouse strains including the C57BL/6 (Twele et al., 2016), which were used in the present study. Differing definitions for HPDs could contribute to this discrepancy. HPDs in our study were defined as series of spikes ≥10 sec and with a frequency >1 Hz in accordance with our previous studies in males. Twele *et al*. applied a more stringent definition requiring a longer duration of ≥20 sec and a higher frequency of 10 – 20 Hz, resembling closely the original definition of HPDs by Riban *et al*. (Riban et al., 2002). Several recent studies, however, define HPDs even more broadly than we do with a minimum duration of 5 sec (Duveau et al., 2016; Zeidler et al., 2018). Akin to Riban *et al*., Twele *et al*. distinguished between HPDs and HVSWs (high-voltage sharp waves) with a duration of 4 – 20 sec and a frequency of 3 – 8 Hz. Owing to these different definitions, the majority of HPDs measured in our study would fall under the definition of HVSWs according to the criteria of Twele *et al*. They described HVSWs as the predominant type of EEG abnormality in females, occurring at a similar rate as in males and progressing over time, which would be comparable to the events measured as HPDs in our analysis. Our observations of an average number of <1 generalized seizure per day are in line with the data of Twele *et al*. Furthermore, they described a faster onset of seizures without a clear seizure-free latent period in the females in contrast to the males showing a latent period of about 2 weeks. In our study we observed a more heterogenous development of female mice after IHKA. Some presented epileptiform activity already in the first recording 1 week after IHKA, which would correspond to an early onset, while others showed a latent period before developing epileptiform activity. Based on the rationale that we generally use mice with >50 sec/h HPDs to test potential antiepileptic interventions because they are considered as having stably established chronic focal epilepsy, we analyzed subgroups. Females with sufficient HPDs (>50 sec/h) at 2 months showed increasing epileptiform activity during the whole post-KA observation period. In contrast, animals that did not develop enough HPDs had a more variable development with a decrease of spike trains and HPDs and occurrence of generalized seizures in some animals.

Moreover, we proved that female IHKA mice replicated neuropathological alterations of human TLE, consistent with previous studies in males (Bouilleret et al., 1999; Riban et al., 2002). Specifically, we observed ipsilateral hippocampal neurodegeneration, mainly in CA1 and less frequently in CA3, accompanied by extensive granule cell dispersion in all animals with HPDs and/or generalized seizures.

While HPDs in male mice after IHKA were described as a model of drug-resistant seizures (Duveau et al., 2016; Riban et al., 2002; Zangrandi et al., 2016), the only previous study testing ASDs in female IHKA mice did not assess HPDs, but so-called seizure-like events with a duration ≥3 sec (Klein et al., 2015). Since drug-responsiveness of HPDs in female IHKA mice has not been addressed yet, we sought to test LTG, OXC and LEV, which are listed among the most frequently prescribed ASDs over the last decade (Hochbaum et al., 2022; Yu et al., 2021). Of each of these ASDs a dose effective in rodent seizure models (Barton et al., 2001; Klitgaard et al., 1998; Luszczki et al., 2003; Luszczki & Czuczwar, 2004) and comparable to the therapeutic range for epilepsy patients as well as a threefold higher dose were tested, consistent with the doses previously applied in our group to evaluate drug-resistance of HPDs in male IHKA mice (Zangrandi et al., 2016). As a positive control we applied the benzodiazepine DZP, which effectively suppressed drug-resistant HPDs in male (Duveau et al., 2016; Riban et al., 2002; Zangrandi et al., 2016) as well as seizure-like events in female IHKA mice (Klein et al., 2015). To account for the previously reported delayed onset of antiseizure activity after i.p. administration in rodents (Castel-Branco et al., 2005; Klitgaard et al., 1998; Löscher, 2007; Luszczki & Czuczwar, 2004; Markowitz et al., 2010), we analyzed a period of 35-94 min after treatment. Inter-individual differences and potential confounding influences of different estrous cycle stages on baseline seizure activity were eliminated by normalizing to the respective pre-treatment period on the same day.

Regarding HPDs, all tested ASDs at the lower dose did not show statistically significant effects, which is in line with the studies in males (Duveau et al., 2016; Zangrandi et al., 2016). Among the tested ASDs only the higher dose of LTG significantly reduced HPDs, while simultaneously increasing spike trains. This shift from HPDs towards spike trains corresponds to a reduction of the duration of individual epileptiform events, while the total number of epileptiform events was not affected. Power spectral analysis using FFT revealed a significant increase in the 1 - 4 Hz frequency band. Interestingly, Duveau *et al*. described in male IHKA mice a biphasic effect of LTG with an increase of HPDs after the same dose of 30 mg/kg, while even higher doses caused a decrease (Duveau et al., 2016). However, the broader definition of HPDs (≥5 sec) in their study incorporates parts of the events that we count as spike trains. Intriguingly, we observed a significant clustering of generalized seizures after LTG, whereas no generalized seizures occurred in the other treatment groups. Seizure aggravation after LTG is known from patients with juvenile myoclonic epilepsy (Gesche et al., 2021), but to our knowledge has not been described in TLE patients. As expected, DZP significantly reduced spike trains and completely abolished HPDs. This was further corroborated by a decrease over a wide range of the power spectrum, with a significant reduction of the 1 – 4 Hz, 4 – 8 Hz, 8 – 13 Hz and 13 – 30 Hz bands as well as the coastline.

Overall, HPDs in female IHKA mice were refractory to established doses of frequently used current ASDs but suppressed by the benzodiazepine DZP, which is in line with the previously reported drug-refractoriness of HPDs in male IHKA mice (Duveau et al., 2016; Riban et al., 2002; Zangrandi et al., 2016) and with the high rate of drug-resistant seizures in TLE patients (Asadi-Pooya et al., 2017; Chipaux et al., 2016; Kuzmanovski et al., 2016; Pohlen et al., 2017). In this regard, female mice after IHKA fulfill the requirements for a model of pharmaco-resistant seizures poorly responding to at least two current ASDs (Stables et al., 2003) and are suitable for the screening of novel ASD candidates.

When developing treatments for patients with drug-resistant seizures, poor water solubility of drug candidates is a frequent problem. Since literature indicated that the highly efficient solvent DMSO may have a confounding influence in epilepsy research, we aimed to clarify its effects on epileptiform activity in IHKA mice of both sexes. Preferably, we sought to identify a concentration that could be used without interfering with the readout. Our results showed that only the highest administered dose of 100% DMSO in 1.5 µl/g (1651 mg/kg) decreased HPDs, suggesting that reducing the DMSO content of a solvent mixture below 30% (495 mg/kg) would be sufficient to minimize confounding effects. This is in accordance with a study in a rat model of electrically induced temporal lobe seizures showing reduced maximal dentate activation only after the highest dose of 1651 mg/kg DMSO (Carletti et al., 2013). In a rat model of genetic absence epilepsy, the contrary effect of increased epileptic activity was observed during later time intervals 30 - 270 min after high doses of DMSO (825 mg/kg and 1651 mg/kg) (Kovács et al., 2011). Our data, in contrast, revealed a rapid onset but short-lasting decrease of epileptiform activity after DMSO 1651 mg/kg compared to saline (**Fehler! Verweisquelle konnte nicht gefunden werden**.) and no significant rebound increase during a later time interval of 35-94 min after treatment (**Fehler! Verweisquelle konnte nicht gefunden werden**.). A potential confounding influence of local painful sensations caused by DMSO injection was excluded by testing DMSO combined with the local anesthetic lidocaine, which did not show statistically significant effects compared to DMSO alone.

In contrast to the observed effect of DMSO in the IHKA mouse model of chronic epilepsy, the threshold of PTZ-induced acute seizures in non-epileptic mice was not affected by DMSO. These discrepancies between different experimental rodent models might be explained by differences in the involved structures and molecular processes.

The potential mechanisms underlying DMSO-induced effects are not clearly understood so far and remain to be elucidated in future studies. Due to its influence on different ion channels (Jacob & de la Torre, 2009; Larsen et al., 1996; Lu & Mattson, 2001; Nakahiro et al., 1992; Santos et al., 2003), DMSO could shift the excitation/inhibition balance and thereby affect neuronal excitability.

In conclusion, we confirmed that female mice after IHKA reproduce essential EEG and morphological features of human TLE patients that were previously modeled in male IHKA mice, in particular drug-resistance of HPDs. Furthermore, we showed that the solvent DMSO caused a significant short-term reduction of HPDs in IHKA mice of both sexes, highlighting that the choice of solvent and appropriate vehicle control is crucial to minimize undesirable misleading effects. Considering the special issues complicating the therapeutic management of women, inclusion of females in the quest for novel treatment strategies is imperative and has in recent years been demanded by funding agencies such as the NIH (Clayton & Collins, 2014). By establishing the female IHKA model we have laid the foundation for future research addressing sex-specific aspects.

## Funding

This work was financed by the Austrian Science Fund (FWF) grants P-30592 and P-30430 (to CS) as well as SPIN W-1206.

## Author contributions

MW planned experiments, performed surgeries and treatments, participated in EEG recordings and analyses, took microscope images, performed statistical analyses and wrote the manuscript. AL programmed the python-based EEG analysis software and power analysis software. AS and BF performed EEG recordings and analyses, brain tissue sections and Nissl stainings. AM contributed to programming the python-based EEG analysis software. CS supervised the study, performed PTZ-seizure threshold experiments and reviewed and edited the manuscript.

## Conflicts of interest

None of the authors has any conflict of interest to disclose.

## Notes

### Competing Interest Statement

The authors have declared no competing interest.

