## Supplemental figures for "Characterization of the intrahippocampal kainic acid model in female mice with a special focus on seizure suppression by antiseizure drugs and DMSO"

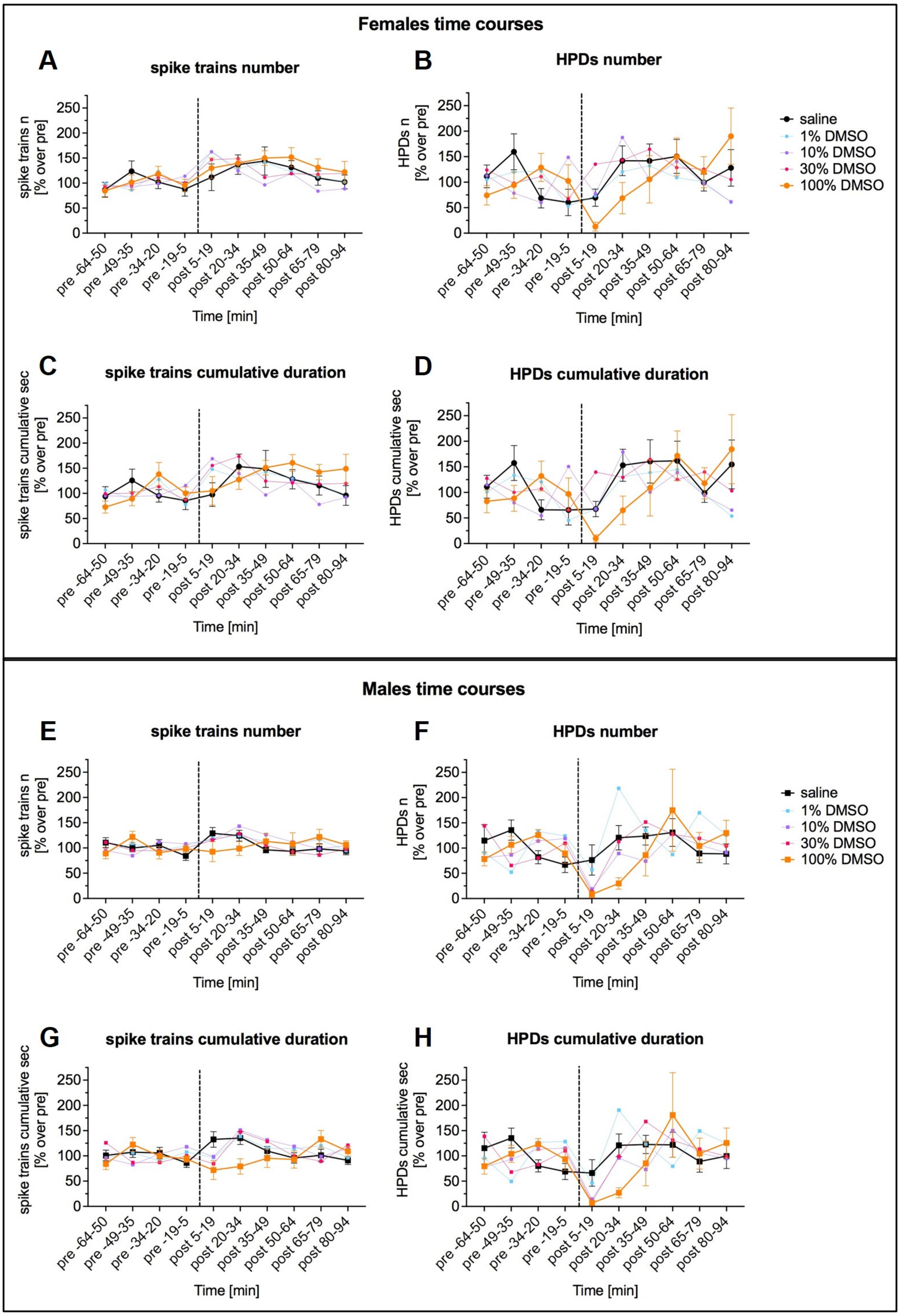

***Supplemental figure 1.*** **Time courses showing the effect of different concentrations of DMSO in female (A-D) and male (E-H) IHKA mice.**
(A, E) Number [% over pre] and (C, G) cumulative duration [% over pre] of spike trains and (B; F) number and (D, H) cumulative duration of HPDs. Data (mean ± SEM for saline and 100% DMSO; for reasons of clarity only mean for 1%, 10% and 30% DMSO) are presented in 15 min bins as % normalized to an average of the pre-treatment period for the 64-5 min before and 5-94 min after the treatment, excluding a handling period of 5 min before and after the treatment. Note the decrease of number and cumulative duration of HPDs in the 5-19 and 20-34 min bins after 100% DMSO compared to saline. Treatment time points are marked by a vertical dashed line. The saline and 100% DMSO curves are highlighted with thicker lines.

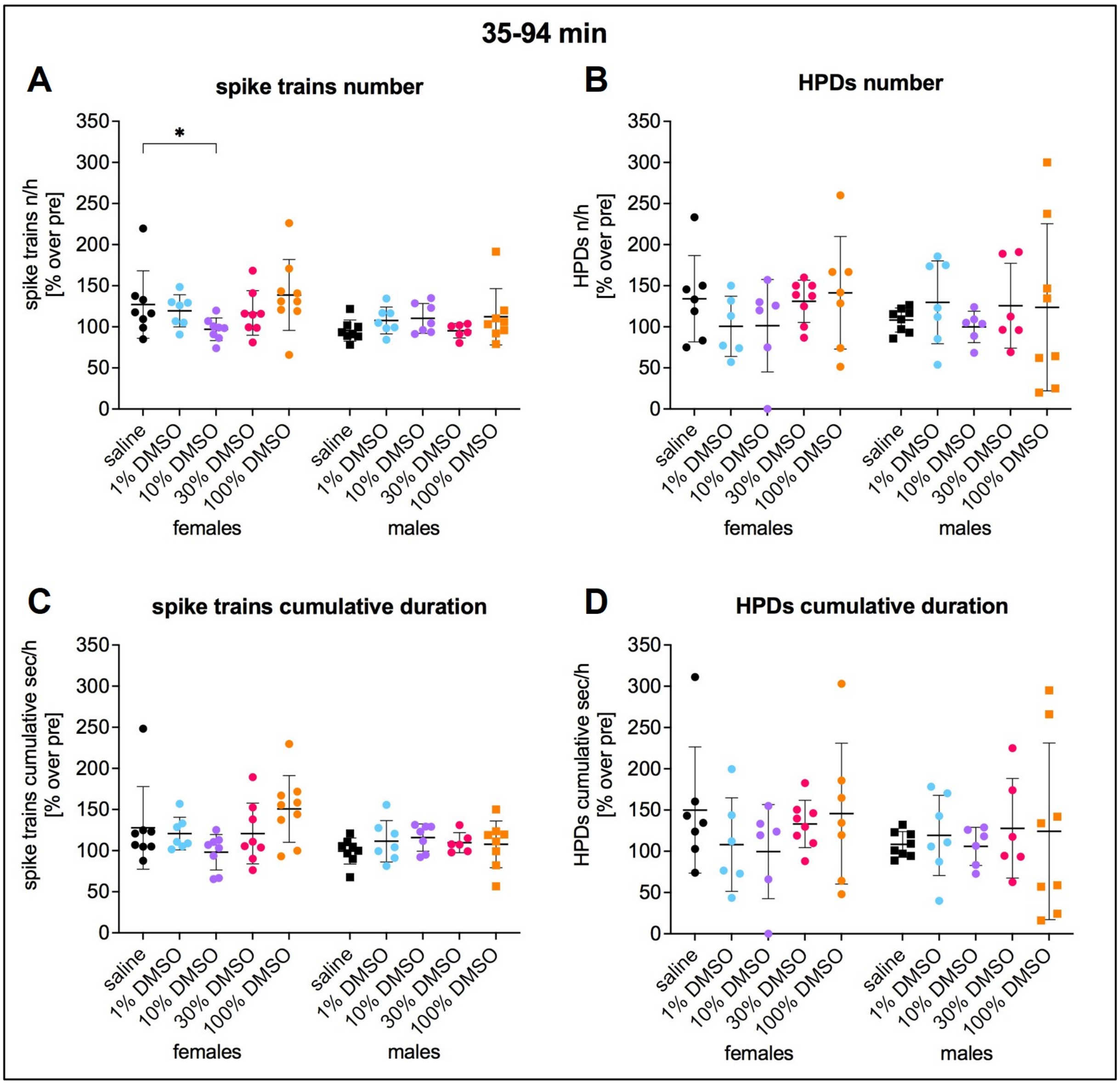


***Supplemental figure 2.*** **Effect of different concentrations of DMSO in IHKA mice in the 35-94 min after treatment.**
(A-D) Effect of 1%, 10%, 30% and 100% DMSO compared to saline in female and male IHKA mice on (A) number [% over pre] and (C) cumulative duration [% over pre] of spike trains and (B) number [% over pre] and (D) cumulative duration [% over pre] of HPDs. No rebound increase was observed in the 35-94 min interval after the initial reduction of spike trains and HPDs following 100% DMSO. Data (females n=9, males n=8) are presented as % normalized to the pretreatment period (mean ± SD) and were analyzed with a 2-way linear mixed model for repeated measures followed by Dunnett's multiple comparisons test. * $p = 0.01-0.05$

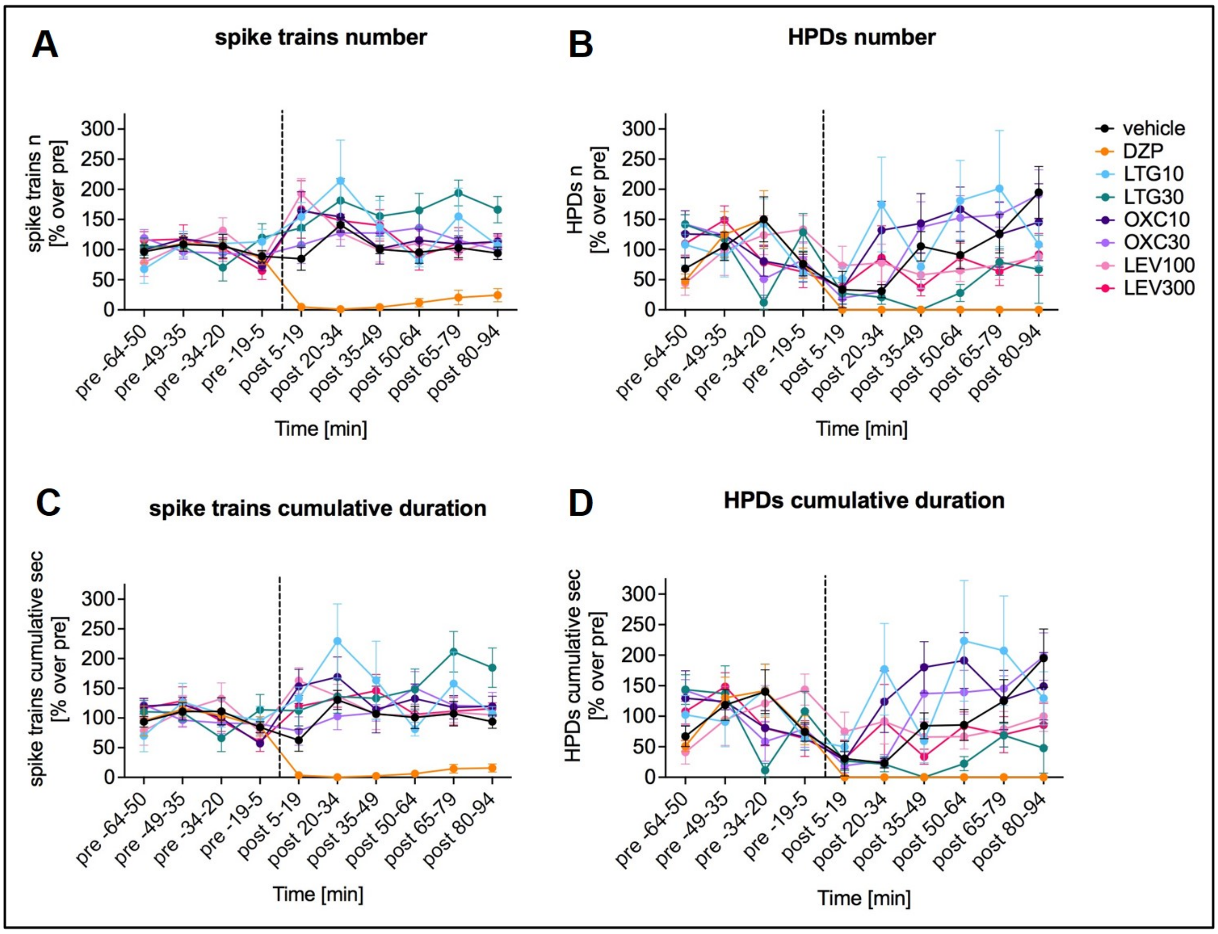


***Supplemental figure 3.*** **Time courses showing the effect of widely prescribed new generation ASDs in female IHKA mice.**

(A) Number [% over pre] and (C) cumulative duration [% over pre] of spike trains and (B) number and (D) cumulative duration of HPDs (B, D). Data (mean ± SEM) are presented in 15 min bins as % normalized to an average of the pretreatment period for the period of 64 – 5 min before and 5 – 94 min after the treatment, excluding a handling period of 5 min before and after the treatment. DZP resulted in a marked decrease of all analyzed parameters in the whole plotted period of 5-94 min after injection. Treatment time points are marked by a vertical dashed line.
